# Haplotype-aware graph indexes

**DOI:** 10.1101/559583

**Authors:** Jouni Sirén, Erik Garrison, Adam M. Novak, Benedict Paten, Richard Durbin

## Abstract

**Motivation:** The variation graph toolkit (VG) represents genetic variation as a graph. Although each path in the graph is a potential haplotype, most paths are nonbiological, unlikely recombinations of true haplotypes.

**Results:** We augment the VG model with haplotype information to identify which paths are more likely to exist in nature. For this purpose, we develop a scalable implementation of the graph extension of the positional Burrows–Wheeler transform (GBWT). We demonstrate the scalability of the new implementation by building a whole-genome index of the 5,008 haplotypes of the 1000 Genomes Project, and an index of all 108,070 TOPMed Freeze 5 chromosome 17 haplotypes. We also develop an algorithm for simplifying variation graphs for k-mer indexing without losing any k-mers in the haplotypes.

**Availability:** Our software is available at https://github.com/vgteam/vg, https://github.com/jltsiren/gbwt, and https://github.com/jltsiren/gcsa2.

**Supplementary information:** Supplementary data are available.

## 1 Introduction

Sequence analysis pipelines often start by mapping the sequence reads to a *reference genome* of the same species. A read aligner first uses a *text index* to find candidate positions for the read. Then it aligns the read to the candidate positions, trying to find the best mapping.

A reference genome that takes the form of a single sequence may represent a new dataset poorly if the sequenced individual diverges substantially at some location. Mapping reads to such a reference can introduce *reference bias* into the subsequent analysis. Richer reference models can help to avoid the bias, but challenges remain in choosing the right model and working with it effectively (The Computational Pan-Genomics Consortium, 2018; Paten *et al.*, 2017).

We can replace the single reference sequence with a collection of haplotypes. Because individual genomes are similar, compressed text indexes can store such collections in very little space (Mäkinen *et al.*, 2010). However, due to this similarity, most reads map equally well to many haplotypes. If the reference model is a simple collection, we cannot tell whether a read maps to the same position in different haplotypes or not.

If the haplotypes are aligned, we can use the alignment to determine whether the mappings are equivalent. Text indexes can also take advantage of the alignment by storing shared substrings only once (Huang *et al.*, 2010). The FM-index of alignment (Na *et al.*, 2016, 2018) goes one step further by collapsing the multiple alignment into a *directed acyclic graph* (DAG), where each node is labeled by a sequence. It indexes the graph and stores some additional information for determining which paths correspond to valid haplotypes.

We can also build a reference graph directly from a reference sequence and a set of variants (Schneeberger *et al.*, 2009). This approach has been used in many tools such as BWBBLE (Huang *et al.*, 2013), GCSA (Sirén *et al.*, 2014), vBWT (Maciuca *et al.*, 2016), the Seven Bridges Graph Pipeline (Rakocevic *et al.*, 2019), Graphtyper (Eggertsson *et al.*, 2017), and VG (Garrison *et al.*, 2018). It will always produce a DAG if structural variants are not considered. Algorithms for working with sequences are often easy to generalize to DAGs. On the other hand, because an acyclic graph imposes a global alignment on the haplotypes, allowing only matches, mismatches, and indels, it cannot represent structural variation such as duplications or inversions adequately.

Assembly graphs such as *de Bruijn graphs* collapse sequences by local similarity instead of global alignment. They are better suited to handling structural variation than DAGs. However, the lack of a global coordinate system limits their usefulness as references.

Graph-based reference models share certain weaknesses. Because they collapse sequences between variants, they represent both the original haplotypes and their *recombinations*: paths that switch between haplotypes. This may cause false positives when a read maps better to an unobserved recombination than to the correct path. Graph regions with many variants in close proximity can give rise to very large numbers of recombinant paths, and be too complex to allow an index to cover all possible paths in the graph. Graph tools try to deal with such regions by, for example, limiting the amount of variation in the graph, artificially simplifying complex regions, and making trade-offs between query performance, index size, and maximum query length.

CHOP (Mokveld *et al.*, 2018) embeds haplotypes into a graph and indexes the corresponding paths. For a given parameter *k*, the graph is transformed into a collection of short strings such that adjacent strings overlap by *k* − 1 characters. Each haplotype can be represented as a sequence of adjacent strings. Any read aligner can be used to map reads to the strings. However, because the aligner sees only short strings, it cannot map long reads or paired-end reads.

The *variation graph toolkit* (VG) (Garrison *et al.*, 2018) works with many kinds of graphs. While some other graph tools use graphs to represent other data types (e.g. aligned sequences, or variants), the graph itself is the primary object in the VG model. A global coordinate system can be provided by designating certain paths as *reference paths*.

VG uses GCSA2 (Sirén, 2017) as its text index. GCSA2 represents a *k*-mer index as a de Bruijn graph and compresses it structurally by merging redundant nodes. VG handles complex graph regions by indexing a *simplified graph*, in which the complex regions are replaced by simpler graph structures, although the final alignment is done in the original graph. The drawback of this approach is that simplification can break paths corresponding to known haplotypes, while leaving paths representing recombinations intact.

In this paper, we augment the VG model with haplotype information. We develop the GBWT, a scalable implementation of the graph extension of the positional Burrows–Wheeler transform (gPBWT) (Durbin, 2014; Novak *et al.*, 2017), to store the haplotypes as paths in the graph. To demonstrate the scalability of the GBWT, we build a whole-genome index for the 1000 Genomes Project (The 1000 Genomes Project Consortium, 2015) haplotypes and a chromosome 17 index for the TOPMed haplotypes. We also describe an algorithm that adds the haplotype paths back to the simplified graph, without reintroducing too much complexity, in order to make the text index more complete.

The main differences from the old gPBWT implementation (Novak *et al.*, 2017) are:

- We use local structures for each node instead of global structures for the graph. The index is smaller and faster and takes better advantage of memory locality.
- The GBWT is implemented as an ordinary text index instead of a special-purpose index for paths. Most FM-index algorithms can be used with it. For example, we can use the GBWT as an FMD-index (Li, 2012) and support *bidirectional search*.
- We have a fast and space-efficient incremental construction algorithm that does not need access to the entire collection of haplotypes at the same time.
- Our implementation can be used independently of VG.

A preliminary version of this paper appeared in WABI 2018. In addition to a more extensive description and discussion, for this paper we have improved the GBWT implementation in the following ways:

- VG now parses the VCF files once and stores the information in a format directly usable by GBWT construction. This makes index construction for the 1000 Genomes Project haplotypes almost three times faster than in the preliminary paper.
- We further demonstrate the scalability of the GBWT by building a chromosome 17 index for the 108,070 TOPMed haplotypes from Freeze 5b, showing that we can build indexes for population cohorts more than an order of magnitude larger than in the original paper.
- We adapt the BWT-merge algorithm (Sirén, 2016) for merging GBWT indexes over the same chromosome. By building indexes for multiple batches in parallel and merging them with the new algorithm, we make GBWT construction for the TOPMed haplotypes several times faster.
- We can now remove paths from a GBWT index.

The haplotype information stored in GBWT can also be used to improve read mapping, and, potentially, variant inference. For example, when we extend the seeds we get from a GCSA2 index, we can restrict the extension to paths corresponding to the haplotypes in the GBWT index. Alternatively, we may use the haplotype information for alignment scoring. We intend to explore these applications in a subsequent paper.

## 2 Methods

### 2.1 Background

#### 2.1.1 Strings and graphs

A *string S*[0, *n* − 1] = *s*_0_ … *s*_*n*−1_ of length |*S*| = *n* is a sequence of *characters* over an *alphabet* Σ = {0, … , *σ* − 1}. *Text* strings *T*[0, *n* − 1] are terminated by an *endmarker T*[*n* − 1] = $ = 0 that does not occur anywhere else in the text. *Substrings* of string *S* are sequences of the form *S*[*i, j*] = *s*_*i*_ … *s*_*j*_. We call substrings of length *k k*-mers and substrings of the type *S*[0, *j*] and *S*[*i*, *n* − 1] *prefixes* and *suffixes*, respectively.

Let *S*[0, *n* − 1] be a string. We define *S.*rank(*i*, *c*) as the number of occurrences of character *c* in the prefix *S*[0, *i* − 1]. We also define *S.*select(*i*, *c*) = max{*j* ≤ *n* | *S.*rank(*j*, *c*) < *i*} as the position of the occurrence of rank *i* > 0. A *bitvector* is a data structure that stores a binary sequence and supports efficient rank/select queries over it.

A *graph G* = (*V*, *E*) consists of a finite set of *nodes V* ⊂ ℕ and a set of *edges E* ⊆ *V* × *V*. We assume that the edges are *directed*: (*u*, *v*) ∈ *E* is an edge *from* node *u to* node *v*. The *indegree* of node *v* is the number of *incoming* edges to *v*, while the *outdegree* is the number of *outgoing* edges from *v*. Let *P* = *v*_0_ … *v*_|*P*|−1_ be a string over the set of nodes *V*. We say that *P* is a *path* in graph *G* = (*V*, *E*), if (*v*_*i*_, *v*_*i*+1_) ∈ *E* for all 0 ≤ *i* < |*P*| − 1.

The VG model (Garrison *et al.*, 2018) is based on *bidirected* graphs, where each node has two *orientations*. We simulate them with directed graphs. We partition the set of nodes *V* into *forward* nodes *V*_*f*_ and *reverse* nodes *V*_*r*_, with *V*_*f*_ ∩ *V*_*r*_ = ∅ and |*V*_*f*_ | = |*V*_*r*_|. We match each forward node *v* ∈ *V*_*f*_ with the corresponding reverse node 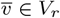, with 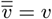 for all *v* ∈ *V*_*f*_. For all nodes *u*, *v* ∈ *V*, we also require that 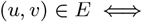 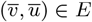.

#### 2.1.2 FM-index

The *suffix array* SA[0, *n* − 1] of text *T*[0, *n* − 1] is an array of pointers to the suffixes of the text in *lexicographic order*. For all *i* < *j*, we have *T*[SA[*i*], *n* − 1] < *T*[SA[*j*], *n* − 1]. The *Burrows*–*Wheeler transform* (BWT) (Burrows and Wheeler, 1994) is a permutation of the text with a similar combinatorial structure. We define it as the permuted string BWT[0, *n* − 1], where BWT[*i*] = *T*[(SA[*i*] − 1) mod *n*]. Let C[*c*] be the number of occurrences of characters *c*′ < *c* in the text. The main operation provided by the BWT is the last-first or *LF-mapping*, which we define as LF(*i*, *c*) = C[*c*] + BWT.rank(*i*, *c*). We use shorthand LF(*i*) for LF(*i*, BWT[*i*]) and note that SA[LF(*i*)] = (SA[*i*] − 1) mod *n*.

Let *X* be a string and let *c* be a character. If *T*′ < *X* for *i* suffixes *T*′ of text *T*, we say that string *X* has *lexicographic rank i* among the suffixes of text *T*. The number of suffixes starting with any character *c*′ < *c* is C[*c*], and the number of suffixes *T*′ < *X* preceded by character *c* is BWT.rank(*i*, *c*). Hence the lexicographic rank of string *cX* is LF(*i*, *c*).

The FM-index (Ferragina and Manzini, 2005) is a text index based on the BWT. Assume that we can compute BWT.rank(*i*, *c*) in *t*_*r*_ time. Further assume that we have stored (*i*, SA[*i*]) for all SA[*i*] divisible by some integer *d* > 0. The FM-index supports the following queries:

- find(*X*): Return the *lexicographic range* [*sp, ep*] of suffixes starting with *pattern X*. If [*sp*_*i*+1_, *ep*_*i*+1_] is the lexicographic range for pattern *X*[*i* + 1, |*X*| − 1], the range for pattern *X*[*i*, |*X*| − 1] is [LF(*sp*_*i*+1_, *X*[*i*]), LF(*ep*_*i*+1_ +1, *X*[*i*])−1]. By extending the pattern backwards, we can support find(*X*) in O(|*X*| · *t*_*r*_) time.
- locate(*sp*, *ep*): Return the list of *occurrences* SA[*sp*, *ep*]. For each position *i* ∈ [*sp*, *ep*], we iterate LF(*i*) until we find a stored pair (LF^*k*^ (*i*), SA[LF^*k*^ (*i*)]). Then SA[*i*] = SA[LF^*k*^ (*i*)] + *k*. Locating each occurrence SA[*i*] takes O(*d* · *t*_*r*_) time.
- extract(*j*, *j*′): Return the substring *T*[*j*, *j*′]. We start from the nearest stored (*i*, SA[*i*]) with SA[*i*] > *j*′ and iterate (*i*, SA[*i*]) ← (LF(*i*), SA[*i*] − 1) until SA[*i*] = *j* + 1. As BWT[*i*] = *T*[SA[*i*] − 1], we extract the substring backwards in O((*d* + *j*′ − *j*) · *t*_*r*_) time.

A generalized FM-index can index *multiple texts T*_0_, … , *T*_*m*−1_. Each text *T*_*j*_ is terminated by a distinct endmarker $_*j*_, where $_*j*_ < $_*j*+1_ for all *j*. As the suffixes of the texts are all distinct, we can sort them unambiguously. In the final BWT, we replace each $_*j*_ with $ in order to reduce alphabet size. The index works as with a single text, except that we cannot compute LF(*i*, $). We also define the *document array* DA as an array of *text identifiers*. If SA[*i*] points to a suffix of text *T*_*j*_, we define DA[*i*] = *j*.

### 2.2 Indexing haplotypes

The positional BWT (Durbin, 2014) and its graph extension (Novak *et al.*, 2017) can be understood as ordinary FM-indexes. We develop the GBWT explicitly from this point of view, making it an FM-index of multiple texts over an integer alphabet *V*. Instead of storing the BWT as a single string, we partition it into substrings BWT_*v*_ corresponding to the most significant character *v* ∈ *V* in the lexicographic ordering. We also use the substrings as the basic blocks for computing rank over the BWT. By storing the substring BWT_*v*_ and rank information in each node *v* ∈ *V*, the resulting index can take advantage of memory locality when the graph has a cache-friendly memory layout.

#### 2.2.1 Positional BWT

Assume that we have *m* haplotype strings *S*_0_, … , *S*_*m*−1_ of equal length over alphabet Σ. At each variant site *i*, character *S*_*j*_ [*i*] tells whether haplotype *j* contains the reference allele (*S*_*j*_ [*i*] = 0) or an alternate allele (*S*_*j*_ [*i*] > 0). Given a pattern *X* and a range of sites [*i*, *i*′], we want to find the haplotypes *S*_*j*_ matching the pattern at the specified sites (*S*_*j*_ [*i*, *i*′] = *X*). Ordinary FM-indexes do not support such queries, as they find all occurrences of the pattern, not just those at a particular position.

The *positional BWT* (PBWT) (Durbin, 2014) is an FM-index that supports *positional queries*. We can interpret it as the FM-index of texts *T*_0_, … , *T*_*m*−1_ such that *T*_*j*_ [*i*] = (*i*, *S*_*j*_ [*i*]) (Gagie *et al.*, 2017). If we want to search for pattern *X* in range [*i*, *i*′], we search for pattern *X*′ = (*i*, *X*[0]) … (*i*′, *X*[|*X*| − 1]) in the FM-index. The texts are over a large alphabet, but their first-order empirical entropy is low. We can encode the BWT using alphabet Σ with a simple model. Assume that SA[*x*] points to a suffix starting with (*i* + 1, *c*). We often know the character from a previous query, and we can determine it using the C array. Then BWT[*x*] = (*i*, *c*′) for a *c*′ ∈ Σ, and we can encode it as *c*′. (Note that we build the rank structure for the original BWT, not the encoded BWT.) When the collection of haplotype strings is repetitive, as it typically is with sufficiently large collections of biological haplotypes, we can compress the PBWT further by run-length encoding the BWT (Mäkinen *et al.*, 2010).

#### 2.2.2 Graph extension

Haplotypes correspond to paths in the VG model. Because chromosome-length phasings are often not available, there may be multiple paths for each haplotype. The *graph extension* of the PBWT (Novak *et al.*, 2017), or gPBWT, generalizes the PBWT to indexing such paths. While the original extension was specific to VG graphs, we present a general version over directed graphs. We call this structure the *Graph BWT* (GBWT), as it both represents arbitrary collections of paths over graphs, and is encoded locally within the graph.

Let *P*_0_, … , *P*_*m*−1_ be paths in graph *G* = (*V*, *E*). We can interpret the paths as strings over alphabet *V*. Assume that 0 ∉ *V*, as we use it as the endmarker. We build an FM-index for the reverse strings. We sort reverse prefixes in lexicographic order, so the LF-mapping traverses edges in the correct direction, and place the endmarker before the string.

The GBWT supports the following variants of the basic FM-index queries:

- find(*X*) returns the lexicographic range of reverse prefixes starting with the reverse pattern (the range of prefixes ending with the pattern).
- locate(*sp*, *ep*) returns the text identifiers DA[*sp*, *ep*]. We do not return text offsets, as the node corresponding to the range [*sp*, *ep*] already provides similar information.
- extract(*j*) returns the path *P*_*j*_. We save memory by not supporting substring extraction.

These queries should be understood as examples of what we can support. Because the GBWT is an FM-index of multiple texts, most algorithms using an FM-index can be adapted to use the GBWT. For example, let *P* = *v*_0_ … *v*_|*P*|−1_ be a path. The *reverse path* of *P* is the path 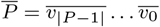 traversing the reverse nodes in the reverse order. If we also index 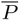 for every path *P*, the GBWT becomes an FMD-index (Li, 2012) that supports bidirectional search.

#### 2.2.3 Records

For each node *v* ∈ *V*, we define the *local alphabet* Σ_*v*_ = {*w* ∈ *V* | (*v*, *w*) ∈ *E*}. We also add $ to Σ_*v*_ if *v* is the last node on a path, and define Σ_$_ as the set of the initial nodes on each path. We partition the BWT into substrings BWT_*v*_ corresponding to the prefixes ending with *v*, and encode each substring BWT_*v*_ using the local alphabet Σ_*v*_. If *w* ∈ Σ_*v*_ is the *k*th character in the local alphabet in sorted order, we encode it as Σ_*v*_ (*w*) = *k*.

We develop a representation based on the following assumptions:

1. Almost all nodes *v* ∈ *V* have a low outdegree, making the local alphabet Σ_*v*_ small. Hence we can afford storing the rank of all *w* ∈ Σ_*v*_ at the start of BWT_*v*_. Decompressing that information every time we access the node does not take too much time either.
2. The number of occurrences of almost all nodes is bounded by the number of haplotypes. As the length of BWT_*v*_ is bounded for almost all *v* ∈ *V*, we can afford scanning it every time we compute rank within it.
3. The collection of paths is repetitive. Run-length encoding compresses the BWT well, reducing both index size and the time required for scanning BWT_*v*_.
4. There exists an integer range [*a, b*] such that the set of nodes *V* is a dense subset of the range. Hence we can afford storing some information for all *i* ∈ [*a, b*] without using too much space.
5. The graph is almost linear and almost topologically sorted. The closer to topological order we can store the nodes, the less space we need for graph topology, and the better we can take advantage of memory locality.

All these are reasonable assumptions for a large set of biological haplotype sequences over a variation graph.

We store a *record* consisting of a *header* and a *body* for each node *v* ∈ *V* and for the endmarker $. For each character *w* ∈ Σ_*v*_ in sorted order, the header stores a pair (*w*, BWT.rank(*v*, *w*)), where BWT.rank(*v*, *w*) is the total number of occurrences of character *w* in all BWT_*v*′_ with *v*′ < *v*. The body run-length encodes BWT_*v*_, representing a run of *ℓ* copies of character *w* as a pair (Σ_*v*_ (*w*), *ℓ*). See Figure 1 for an example.

**Fig. 1.**
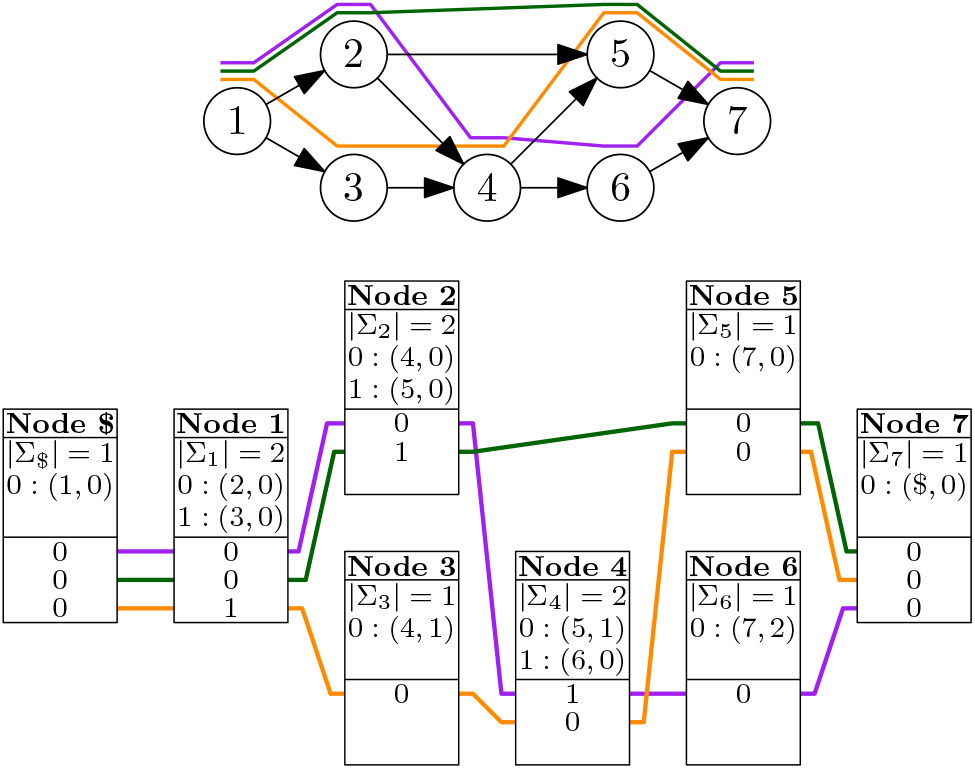
Top: A graph with three paths (colored lines). Bottom: The GBWT of the paths, with correspondingly colored lines connecting the paths’ entries in each node’s record.

Because the BWT is a set of records, we use node/offset pairs as *positions*. Pair (*v*, *i*) refers to offset BWT[C[*v*] + *i*] = BWT_*v*_ [*i*]. We define rank queries over positions as

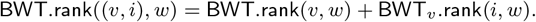

Similarly, we define LF((*v*, *i*), *w*) = (*w*, BWT.rank((*v*, *i*), *w*)) and use it in place of ordinary LF-mapping in the FM-index (see Section 2.1.2).

The FM-index is based on iterating LF-mapping. Because LF-mapping in a standard BWT tends to jump randomly around the BWT, this can be a significant bottleneck. The GBWT achieves better memory locality, if we store the records for adjacent nodes close to each other. When we iterate LF-mapping over a path in the graph, we traverse adjacent memory regions.

As a run-length encoded FM-index, the GBWT supports the *fast* locate() *algorithm* (Mäkinen *et al.*, 2010). The *direct algorithm*, as described in Section 2.1.2, locates each position *i* ∈ [*sp*, *ep*] separately. If we instead process the entire range at once, advancing every position by one step of LF-mapping at the same time, we achieve better memory locality. We can also compute LF-mapping for an entire run BWT_*v*_ [*x*, *y*] = *w*^*y*+1−*x*^ in the same time as for a single position *i* ∈ [*x*, *y*].

#### 2.2.4 GBWT encodings

We have two representations for the GBWT. The *dynamic GBWT* is a representation of the GBWT optimised for index construction, where speed is more important than size. The *compressed GBWT* balances query performance with index size. We use it when the set of haplotypes is fixed and for storing the index on disk. See Supplement 1 for further details.

### 2.3 GBWT construction

The assumptions in Section 2.2.3 make the GBWT easier to build than an ordinary FM-index. *Inserting* new texts into the collection updates adjacent records, just like searching traverses adjacent records. Because the local alphabet is small, because the number of occurrences of each character is limited, and because run-length encoding compresses the BWT well, records tend to be small. Hence we can afford *rebuilding* a record each time we update it.

On the other hand, the GBWT is harder to build than the PBWT. In the PBWT, all strings are of the same length and have the same variant site at the same position. Hence we can build the final record for a site in a single step. In the GBWT, indels in the haplotypes become indels on the haplotype paths, and hence we have to update the same record multiple times. We also have to buffer the strings instead of indexing them as we generate them.

#### 2.3.1 Construction algorithms

Our basic GBWT construction algorithm is similar to RopeBWT2 (Li, 2014). We have a dynamic FM-index (Chan *et al.*, 2007) and insert multiple texts into the index in a single *batch* using the BCR algorithm (Bauer *et al.*, 2013). The algorithm is sequential and hence not suitable for large datasets. Parallelizing it is difficult, because the algorithm interleaves queries with index updates.

When the basic algorithm is too slow, we partition the dataset into *superbatches* and build a separate GBWT index for each superbatch. We then merge the indexes using the BWT-merge algorithm (Sirén, 2016). The merging algorithm can also be reversed to remove texts from the index. This can be useful, if we want to remove a sample from the dataset without having to rebuild the entire index.

Indexes for different chromosomes can be merged quickly with a simple algorithm. Because each chromosome uses different node identifiers, we can simply reuse the existing records in the merged index.

See Supplement 2 for further details on the construction algorithms.

#### 2.3.2 Construction in VG

We provide support in VG to construct a GBWT from a VCF file (Danecek *et al.*, 2011) with phasing information. The construction parses the VCF file, determines the path corresponding to each allele, and builds haplotype paths from the allele paths. Because we need two layers of buffering, we process the VCF file in batches of *s* samples (default 200) in order to save memory. Heuristics are required to deal with situations where VCF sites overlap and paths may not be well defined.

The GBWT stores texts with integer identifiers. A *metadata* layer maps the identifiers to structured names. While the metadata supports genomes with arbitrary ploidy, our current VCF parsing code does not take full advantage of it. The parser expects a diploid genome, where some regions may be haploid.

As repeated VCF parsing can be slower than GBWT construction, we start by parsing the VCF file and storing the information in a directly usable format in a number of files. The *main file* contains the reference path and the paths corresponding to each allele of each variant. For each batch, we create a *phasing file* containing the run-length encoded phasing information for the corresponding samples. The total size of the files is usually comparable to a compressed VCF file. If the VCF file contains interleaved diploid and haploid samples (e.g. female and male samples for chromosome X), the files can be several times larger. After the files have been written, we can continue the construction in VG or use separate GBWT construction tools.

We generate a path for each haplotype in the current batch. At each variant site and for every haplotype, we first append reference nodes until the site. Then we check whether the reference coordinates of the site overlap with the path we have already generated. If there is an overlap, we try to resolve it by removing reference nodes from the generated path or by skipping reference nodes on the path corresponding to the allele at the current site. If we cannot resolve the overlap, we can treat it as a *phase break* and start a new path. Alternatively, we can replace the alternate allele with the reference allele. Finally we append the path corresponding to the allele to the end of the path we are generating.

When we have finished the haplotype or there is a phase break, we insert the path *P* and its reverse 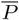 into the GBWT *construction buffer*. Once the buffer is full (the default size is 100 million nodes), we launch a background thread to insert the buffer into the index.

### 2.4 Haplotype-aware graph simplification

VG uses a series of pruning heuristics to simplify graphs for *k*-mer indexing. First it removes edges used by *k*-mers that make too many edge choices (e.g. more than 3 choices in a 24-mer). Edges with no alternatives are not deleted, as there is no choice in taking them. Then it deletes connected components with too little sequence (e.g. less than 33 bases). Finally, if the graph contains reference paths, it may add them back to the pruned graph.

Heuristic pruning often breaks paths taken by known haplotypes. This may cause errors in read mapping, if we cannot find candidate positions for a read in the correct graph region. On the other hand, indexing too many recombinations may increase the number of false positives. Hence we would like to prune recombinations while leaving the haplotypes intact.

We describe an algorithm that *unfolds* the haplotype paths in pruned regions, restoring support for them in the graph, and duplicating nodes when necessary. Our algorithm works with any pruning algorithm that removes nodes from the graph. See Figure 2 for an example of the algorithm in action. We work with bidirected VG graphs, unless otherwise noted. Reference paths can also be unfolded with a similar algorithm.

**Fig. 2.**
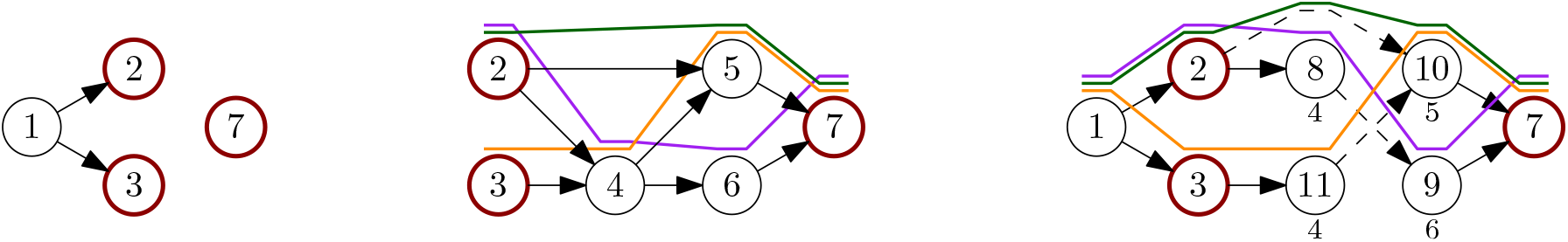
Unfolding the paths in the graph in Figure 1. Border nodes have been highlighted. Left: The graph after removing nodes 4, 5, and 6. Center: Complement graph. The maximal paths are (2, 4 | 6, 7), (2 | 5, 7), and (3, 4 | 5, 7), with the bar splitting a path into a prefix and a suffix. Right: Unfolded graph. Dashed edges cross from prefixes to suffixes. Duplicated nodes have the original ids below the node.

Let *G*_*i*_ = (*V*_*i*_, *E*_*i*_) be the graph induced by GBWT paths and *G*_*p*_ = (*V*_*p*_, *E*_*p*_) be a pruned graph. We build a *complement graph* induced by edges *E*_*i*_ \ *E*_*p*_ and consider each connected component *G*_*c*_ = (*V*_*c*_, *E*_*c*_) in it separately. The set *V*_*b*_ = *V*_*c*_ ∩ *V*_*p*_ is the *border* of the component, as the nodes exist both in the component and in the pruned graph. Nodes in the set *V*_*c*_ \ *V*_*b*_ are *internal nodes*.

Each connected component *G*_*c*_ represents a graph region that was removed from the original graph. We build an unfolded component consisting of the paths in *G*_*c*_ supported by GBWT paths and insert it into the pruned graph *G*_*p*_. We achieve this by duplicating the internal nodes that would otherwise cause recombinations.

In order to build the unfolded component, we must find all *maximal* paths *P* of length |*P*| ≥ 2 supported by GBWT paths in the component. A path starting from a border node is maximal if it reaches the border again or cannot be extended any further. GBWT paths consisting entirely of internal nodes of the component are also maximal.

Let *v* be a GBWT node and vg(*v*) ∈ *V*_*c*_ the corresponding VG node. If vg(*v*) is a border node, we create a *search state* (*v*, find(*v*)) consisting of a pattern and a range. For internal nodes, we create search state (*v*, find($*v*)). Then, for each search state (*X*,[*sp*, *ep*]), with *x* = *X*[|*X*| − 1]:

1. If |*X*| ≥ 2 and the last node vg(*x*) is a border node, we stop the search for this state. If vg(*X*[0]) is also a border node, *X* is a maximal path, and we output it.
2. We try to extend the search with all GBWT nodes *v* corresponding to the successors *u* ∈ *V*_*c*_ of vg(*x*), taking the orientation of *v* from the VG edge. If [*sp*′, *ep*′] = [LF((*x*, *sp*), *v*), LF((*x*, *ep* + 1), *v*) − 1] ≠ ∅, we create a new state (*Xv*, [*sp*′, *ep*′]).
3. If no extension was successful and |*X*| ≥ 2, path *X* is maximal, and we output it.

Let *P* be a maximal path we output. If *P* is not a border-to-border path, we try to extend the lexicographically smaller of *P* and 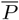 with reference paths, replacing *P* with the extended path. To avoid having the same path in both orientations, we replace each path *P* with the smaller of *P* and 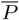.

We could create new duplicates of all internal nodes on *P* and insert the path into *G*_*p*_, but this would create too much nondeterminism for GCSA2.^1^ Instead, we split each path into a prefix and a suffix of equal length and build a trie of the prefixes and a trie of the reverse suffixes. Every edge in the tries becomes a node in the unfolded component.

Let *v* be the label of a trie edge starting from the root. If vg(*v*) is a border node, it already exists in *G*_*p*_. Otherwise we add a new duplicate of vg(*v*). Now let *v* and *v*′ be the labels of two successive trie edges, and let *u* be the VG node we used for *v*. We create a new duplicate *u*′ of vg(*v*′) and add node *u*′ to *G*_*p*_. We also add edge (*u*, *u*′) or (*u*′, *u*), depending on whether we are in a prefix or a suffix. Finally, if we used VG node *u* for the end of a prefix and VG node *u*′ for the start of the corresponding suffix, we add edge (*u*, *u*′) to *G*_*p*_.

After we have handled all components, the simplified graph *G*_*p*_ contains all GBWT paths. The GCSA2 index of *G*_*p*_ contains all *k*-mers (e.g. 256-mers) in the haplotypes. This allows us to prune the graph more aggressively, removing more *k*-mers corresponding to recombinations. In order to map reads to the original graph *G* = (*V*, *E*) instead of the simplified graph *G*_*p*_, we replace the node identifiers *v* ∈ *V*_*p*_ in the GCSA2 index with the original identifiers *v*′ ∈ *V*.

## 3 Results

We have implemented the GBWT in C++ using the SDSL library (Gog *et al.*, 2014). The following experiments were done using VG v1.12.1 with GCSA2 v1.2 and a prerelease version of GBWT v0.8. All code was compiled using GCC 7.3. We used a single Amazon EC2 i3.8xlarge instance with 16 physical (32 logical) cores of an Intel Xeon E5 2686 v4 and 244 GiB^2^ of memory. The system was running Ubuntu 18.04 with Linux kernel 4.15.0. Temporary files were stored on a local RAID 0 volume consisting of four 1.9 TB SSDs.

In the following, we discuss GBWT construction benchmarks and experiments with haplotype-aware graph simplification. Supplement 3 contains low-level query benchmarks that show how the GBWT can take advantage of memory locality.

### 3.1 Datasets

We built VG graphs for two datasets: *1000 Genomes Project* (1000GP) final phase (The 1000 Genomes Project Consortium, 2015) whole-genome haplotypes and *Trans-Omics for Precision Medicine* (TOPMed) Freeze 5b haplotypes for chromosome 17. The 1000GP graphs were built relative to the GRCh37 human reference genome, while the TOPMed graph used the GRCh38 reference. As the 1000GP haplotypes have issues with overlapping variants, we generated both *short paths* by creating phase breaks at unresolvable overlaps, and *long paths* by ignoring the variants that caused such overlaps. The TOPMed haplotypes had only 0.29 unresolvable overlaps per haplotype on the average, so we generated only long paths.

See Table 1 for further details on the datasets and Supplement 4 for a comparison of input (reference, VCF) and output (graph, GBWT) sizes. We have also included the 1000GP chromosome 17 for comparison. The TOPMed graph was built with the old VG maximal node size default of 1,000 bp, while the 1000GP graphs use the new 32 bp default. The effect of this difference is negligible: the average distance between variants in the TOPMed graph is only 6.5 bp.

**Table 1.** Datasets and direct GBWT construction. The name of each dataset is a combination of source, chromosome, and path length (S for short paths with phase breaks, L for long chromosome-length paths). For each dataset, we give the number of samples and variants, as well as the number of nodes, average/maximum outdegree, number of paths (including reverse paths), and the total length of the paths in the graph. For construction, we give batch size in samples, buffer size in millions of nodes, interval for stored text identifiers, and wall-clock time for index construction in hours or days. We also report GBWT size, space used by text identifiers, and total index size. M, G, and T suffixes indicate millions, billions, and trillions, respectively.

| Dataset |  |  | Graph |  |  |  | Construction |  |  |  | Index |  |  |
| --- | --- | --- | --- | --- | --- | --- | --- | --- | --- | --- | --- | --- | --- |
| Name | Samples | Variants | Nodes | Outdegree | Paths | Length | Batch | Buffer | Interval | Time | GBWT | Text IDs | Total |
| 1000GP-all-S | 2,504 | 84.7 M | 612 M | 1.3 / 13 | 50.6 M | 2.19 T | 200 | 100 M | 1,024 | 12 h | 8.74 GiB | 9.90 GiB | 18.6 GiB |
| 1000GP-all-L | 2,504 | 84.7 M | 612 M | 1.3 / 13 | 240,232 | 2.19 T | 100 | 200 M | 1,024 | 17 h | 8.43 GiB | 8.17 GiB | 16.6 GiB |
| 1000GP-17-S | 2,504 | 2.33 M | 16.6 M | 1.3 / 7 | 1.67 M | 60.1 G | 200 | 100 M | 1,024 | 3.7 h | 258 MiB | 242 MiB | 500 MiB |
| 1000GP-17-L | 2,504 | 2.33 M | 16.6 M | 1.3 / 7 | 10,016 | 60.1 G | 100 | 200 M | 1,024 | 4.2 h | 252 MiB | 193 MiB | 444 MiB |
| TOPMed-17-L | 54,035 | 12.9 M | 67.6 M | 1.3 / 11 | 216,140 | 4.66 T | 200 | 1000 M | 16,384 | 25 d | 1.13 GiB | 1.03 GiB | 2.16 GiB |

### 3.2 Index construction

For 1000GP, we first built separate indexes for each chromosome, running 14 jobs in parallel. We used the default construction parameters with short paths. For long paths, we used a larger buffer size (to speed up the construction) and a smaller batch size (to save memory). The number of parallel jobs was determined by memory usage (almost 1 GiB for each 10 Mbp) and the number of CPU cores (2 threads per job). We ordered the jobs *X*, 1, … , 22, *Y*, as large chromosomes take longer to finish. The last job to finish was chromosome 2, which determined the total construction time (11.8 h and 16.3 h for short and long paths, respectively). Merging the single-chromosome indexes into a whole-genome index took 12 minutes and 38 GiB memory for short paths and 12 minutes and 33 GiB for long paths. See Table 1 for further details.

Because the direct construction algorithm is sequential, it is too slow for building the TOPMed index (see Table 1). We used the BWT-merge algorithm for faster parallel construction. Parsing the VCF file and writing the phasing information in batches of 100 samples took 42 hours and 39 GiB memory. We then grouped the batches into 22 superbatches of 2500 samples each and built GBWT indexes for the superbatches in parallel. With a buffer size of 500 million, the construction took 15 hours and 33 GiB memory for each of the first 21 superbatches and 9 hours and 21 GiB for the partially-filled last one. We had enough memory to run 7 jobs in parallel, so the total wall-clock time for indexing the superbatches was 54 hours.

We merged the superbatch indexes in 46 hours and 102 GiB memory. The total construction time was 5.9 days or 4.2 times less than with the direct construction algorithm. By building all superbatch indexes in parallel, the construction time can be reduced further to 4.3 days, out of which 1.7 days is spent for VCF parsing. See Supplement 2 for further details.

The size of the compressed 1000GP whole-genome GBWT was 18.6 GiB with short paths and 16.6 GiB with long paths. Roughly half of this was for the GBWT itself and half for the stored text identifiers. See Table 1 for further details. The decompressed endmarker (see Supplement 1) adds another 386 MiB with short paths and 1.83 MiB with long paths. The TOPMed chromosome 17 index took 2.16 GiB, which was also split roughly in half between the GBWT and the text identifiers. It was 5.0 times larger than the corresponding 1000GP index with long paths, while containing 22 times more samples over 5.5 times more variants. The dynamic indexes are roughly 10 times larger than the corresponding compressed indexes. Their exact sizes are not well-defined due to a large number of memory allocations and unused space in the arrays.

All the assumptions in Section 2.2.3 were valid for our datasets:

1. All nodes except the endmarker have low outdegrees.
2. As the graphs are acyclic, no path can visit the same node twice. When phase breaks or overlapping variants break a haplotype into multiple paths, there will be some overlap between the paths.
3. The 1000GP indexes take 0.03 to 0.04 bits per character and the TOPMed index takes 0.002 bits per character, excluding the text identifiers.
4. VG construction tries to avoid leaving gaps between node identifiers.
5. The VG graphs built from a VCF file are almost in topological order.

### 3.3 Haplotype-aware graphs

VG originally used 128-mer GCSA2 indexes. When pruning the graph for indexing, we could allow 4 edge choices in a 16-mer. Such indexes were unsatisfactory, however, because they could not map exactly matching 150 bp reads in one piece. When we double the order of the index to 256-mers, we need more aggressive pruning to avoid exponential growth during index construction. By default, we now double the distance between edge choices, allowing only 3 edge choices in a 24-mer.

Pruning removes *k*-mers corresponding to both true haplotypes and their recombinations. To determine the effect of more aggressive pruning, we built several whole-genome GCSA2 indexes for the 1000GP graph and compared their *k*-mer contents. In the following, graph pruned-*k* has been pruned with the parameters for a *k*-mer index, and the reference paths have been restored afterwards. Similarly, unfolded-*k* is a graph where the haplotype paths and reference paths have been unfolded after pruning, using the GBWT index with long paths. See Table 2 for the results.

**Table 2.**
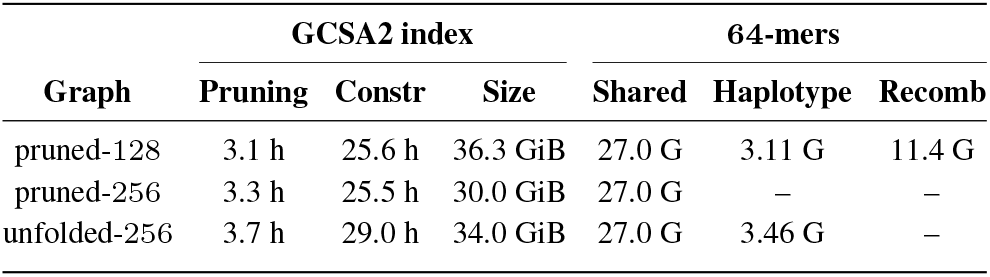
GCSA2 indexes for simplified 1000GP graphs. For each graph, we give the pruning time in hours, construction time in hours, and index size in GiB. We also show a comparison of unique 64-mer content vs. pruned-256, including the number of 64-mers shared with pruned-256, the number of additional (real) haplotype 64-mers over pruned-256, and the number of additional (spurious) recombination 64-mers over pruned-256. The G suffix indicates billions.

| Graph | GCSA2 index |  |  | 64-mers |  |  |
| --- | --- | --- | --- | --- | --- | --- |
|  | Pruning | Constr | Size | Shared | Haplotype | Recomb |
| pruned-128 | 3.1 h | 25.6 h | 36.3 GiB | 27.0 G | 3.11 G | 11.4 G |
| pruned-256 | 3.3 h | 25.5 h | 30.0 GiB | 27.0 G | – | – |
| unfolded-256 | 3.7 h | 29.0 h | 34.0 GiB | 27.0 G | 3.46 G | – |

There are 27.0 billion 64-mers in the pruned-256 index. Unfolding the haplotype paths adds 3.46 billion 64-mers to the unfolded-256 index. While most of these additional *k*-mers correspond to haplotypes, some of them may be recombinations covering multiple complex regions. The pruned-128 index contains 90 % of the haplotype *k*-mers that were missing from pruned-256, but it also adds 11.4 billion 64-mers arising from recombinations of the haplotypes. Overall, by switching from the old pruned-128 index to the new unfolded-256 index, we add the last missing haplotype *k*-mers to the index while getting rid of a large number of recombination *k*-mers, and supporting direct matches of sequencing reads up to 256 bp.

## 4 Discussion

We have developed the GBWT, a scalable implementation of the graph extension of the PBWT. The earlier gPBWT implementation used 9.3 hours and 278 GiB of memory for indexing the 1000GP chromosome 22 using a single thread (Novak *et al.*, 2017). In comparison, our implementation takes 2.0 hours and 4.0 GiB (short paths) or 1.9 hours and 5.9 GiB (long paths) using two threads. We also reduced the final index size from 321 MiB to 134 MiB (short paths) or 131 MiB (long paths) without text identifiers. By running multiple jobs in parallel, we were able to build a whole-genome index in 12 hours (short paths) or 17 hours (long paths) on a single system.

Contemporary sequencing projects are sequencing in excess of 100,000 diploid genomes. Our aim is to scale the GBWT to allow working with such large collections, providing a compressed, indexed and searchable representation that should fit into the memory of a single server. Potential applications in genome inference and imputation, as well as for powering population genomic queries, are myriad. For example, we are exploring using the GBWT for additionally scoring read mappings by the number of recombinations of the underlying haplotypes they induce, using the model described by Rosen *et al.* (2017).

Our experiments with the TOPMed dataset suggest we are almost there. We can build a 2.16 GiB chromosome 17 index for 54,035 diploid samples in 4.3 days. Extrapolating from this, it should take 13–14 days and 76,000 CPU hours to build an 80–90 GiB whole-genome index. See Table 3 for further details. While GBWT merging uses more CPU hours than the other phases, GBWT construction for superbatches requires 5 times more memory per CPU core, so its actual cost may be higher.

**Table 3.** Estimated resource usage of whole-genome TOPMed GBWT construction. For each phase (VCF parsing, GBWT construction for superbatches, GBWT merging), we give the number of jobs per chromosome, number of CPU cores per job, time (in hours or days) and memory usage (in GiB) per job for chromosome 17 (measured) and chromosome 2 (estimated), and estimated CPU hours for building a whole-genome index.

| Phase | Jobs | Cores | Chr 17 |  | Chr 2 |  | CPU hours |
| --- | --- | --- | --- | --- | --- | --- | --- |
|  |  |  | Time | Memory | Time | Memory |  |
| Parsing | 1 | 1 | 42 h | 39 GiB | 5–6 d | 128 GiB | 1,500 |
| Construction | 22 | 2 | 15 h | 33 GiB | 2 d | 100 GiB | 23,000 |
| Merging | 1 | 32 | 46 h | 102 GiB | 5–6 d | 320 GiB | 51,500 |

We can probably improve the resource usage with a better choice of batch/superbatch sizes and other construction parameters. For wall-clock time, the main bottleneck is the sequential VCF parsing, which takes 40 % of the total construction time. Improvements to this may involve integrating the parsing into VG graph construction, parsing multiple graph regions in parallel, or switching to a more efficient input format.

Storing the text identifiers for locate() queries is another bottleneck. When the number of samples increases, the product of locate() time and the space taken by the stored text identifiers increases linearly. In the 1000GP dataset, storing the identifiers at one out of 1,024 positions takes roughly as much space as the GBWT itself. With the TOPMed dataset, we achieve similar proportions by storing the identifiers at one out of 16,384 positions, making locate() 16 to 22 times slower. There is a theoretical proposal for supporting fast locate() queries in space proportional to the size of the run-length encoded BWT (Gagie *et al.*, 2018). While there has been some progress in building the proposed index for large datasets (Kuhnle *et al.*, 2019), scaling it up to TOPMed scale is still an open problem.

We used the haplotype information in the GBWT to simplify VG graphs for *k*-mer indexing. This allowed us to prune the *k*-mers corresponding to recombinations more aggressively, while still having all *k*-mers from the haplotypes in the index. CHOP, the other haplotype-aware graph indexing approach, can only use short-range haplotype information in read mapping. Because VG graphs are connected, we can use the long-range information in the GBWT for mapping long reads and paired-end reads. We will investigate this in a subsequent paper.

## Funding

This work was supported by the National Institutes of Health [5U41HG007234 and 1U01HL137183-01 to JS, AN, and BP]; the Wellcome Trust [WT206194 to JS, EG, and RD; WT207492 to RD]; and the W. M. Keck Foundation [DT06172015 to JS, AN, and BP].

## Footnotes

1 If VG node *v* has predecessors *u* and *u*′ with identical labels, *k*-mers starting from *u* and *u*′ and passing through *v* cannot be distinguished. GCSA2 construction has to extend these *k*-mers until the order of the index (e.g. *k* = 256), which may increase the size of the temporary files significantly.

2 Sizes measured in MiB, GiB, and TiB are based on 1,024-byte kibibytes. Sizes measured in MB, GB, and TB are based on 1,000-byte kilobytes.

## Notes

#### Summary of Updates

Better overview of the GBWT index in Section 2.2. Text clarifications in multiple sections and supplements.

https://github.com/vgteam/vg

https://github.com/jltsiren/gbwt

https://github.com/jltsiren/gcsa2

