## Supplementary Material for "Haplotype-aware graph indexes"

May 29, 2019

#### 1 GBWT encodings

##### 1.1 Dynamic GBWT

The *dynamic GBWT* is a representation of the GBWT optimised for index construction, where speed is more important than size. We have an array of fixed-size records for characters \$ and $v \in [a, b]$, including character values $v \notin V$. The record for the endmarker is at array position 0, while the record for node $v$ is at array position $v - (a - 1)$.

For each node $v \in V$, the corresponding record contains four pointers to arrays: header, body, incoming edges, and text identifiers. For each incoming edge $(u, v) \in E$, the incoming edges array stores a pair $(u, \mathsf{BWT}_u.\mathsf{rank}(|\mathsf{BWT}_u|, v))$, recording the number of paths crossing from $u$ to $v$.

Let $\mathsf{SA}_v$ and $\mathsf{DA}_v$ be the parts of $\mathsf{SA}$ and $\mathsf{DA}$ corresponding to $\mathsf{BWT}_v$. The text identifiers array for node $v$ stores, in sorted order, pairs $(i, \mathsf{DA}_v[i])$ for which $\mathsf{SA}_v[i]$ points to either the last node on a path or a path position divisible by $d > 0$. These pairs are used for $\mathsf{locate}()$ queries, like stored $\mathsf{SA}$ pointers in an ordinary FM-index.

##### 1.2 Compressed GBWT

The *compressed GBWT* balances query performance with index size. We use it when the set of haplotypes is fixed and for storing the index on disk. Each record is a byte array. We encode integers as sequences of bytes, where the lower 7 bits contain data and the high bit tells whether the encoding continues. The header starts with the local alphabet size $|\Sigma_v|$. We encode the outgoing edges $(w_i, \mathsf{BWT}.\mathsf{rank}(v, w_i))$ *differentially*, replacing $w_i$ with $w_i - w_{i-1}$. If the local alphabet is large, each run $(k, \ell)$ in the body is encoded as an integer pair. Otherwise we encode $k$ and as much of $\ell$ as possible in the first byte, and continue with the remaining run length in subsequent bytes.

We concatenate all records and mark their starting positions in a sparse bitvector [5] $B_V$. The records can be accessed with $\mathsf{select}$ queries on the bitvector. If the record for node $v$ is at array position $i$ in the dynamic GBWT (see Supplement 1.1), its encoding starts at position $B_V.\mathsf{select}(i + 1, 1)$ in the concatenated byte array. If array position $i$ does not correspond to any node, the record uses one byte to indicate that the local alphabet size is 0.

Each compressed record must be decompressed sequentially. As the stored text identifiers tend to cluster in certain nodes, storing them in records would make these records large and slow to decompress. Instead, we use a global structure for the text identifiers. The structure consists of three bitvectors and an array of identifiers:

- Uncompressed bitvector $B_s$ marks the records with stored identifiers. If the $i$th record contains identifiers, we set $B_s[i] = 1$. This allows us to skip checking the identifiers in most records when iterating $\mathsf{LF}()$.

- Sparse bitvector $B_r$ is defined over the concatenated offset ranges of the records with stored identifiers. If $B_s[i] = 1$, the range for the record starts at $B_r.\mathsf{select}(B_s.\mathsf{rank}(i, 1) + 1, 1)$.

- Sparse bitvector $B_o$ covers the same range as $B_r$. If $B_o[i + j] = 1$ and the range for the record starts at $B_o[i]$, we have an identifier for offset $j$ at array position $B_o.\mathsf{rank}(i + j, 1)$.

The endmarker record storing $\mathsf{BWT}_\$$ can be very large, if there are many texts in the collection or if the local alphabet $\Sigma_\$$ is large. Because accessing large records is expensive in the compressed GBWT, we decompress the endmarker record for faster access. As the endmarker is mostly used for extracting entire texts, we decompress $\mathsf{BWT}_\$$ into an array $\mathsf{LF}_\$$ such that $\mathsf{LF}_\$[i] = \mathsf{LF}((\$, i), \mathsf{BWT}_\$[i])$. This array takes 8 bytes per text, as long as the number of nodes and the size of the largest $\mathsf{BWT}_v$ are both less than $2^{32}$.

### 2 GBWT construction

#### 2.1 Direct construction

The following algorithm [3] updates the BWT of text $T$ to be the BWT of text $cT$, where $c$ is a character, forming the basis of many incremental BWT construction algorithms:

1. Find the offset $i$ where $\mathsf{BWT}[i] = \$$ and replace the endmarker with character $c$.

2. Compute $i' = \mathsf{LF}(i, c)$ and insert a new endmarker between offsets $i' - 1$ and $i'$.

If we have a BWT for $m$ texts, we can insert a new empty text by inserting an endmarker between offsets $m - 1$ and $m$. By iterating the above algorithm, we can then insert the actual text. If we have a *dynamic FM-index* [2], this can be quite efficient in practice.

The BCR algorithm [1] builds a BWT for $m$ texts. It starts with the BWT for $m$ empty texts and then extends each text by one character in each step. Originally intended for indexing short reads, the BCR algorithm is also used for PBWT construction.

Our GBWT construction algorithm is similar to RopeBWT2 [4]. We have a dynamic GBWT and insert multiple texts into the index in a single *batch* using the BCR algorithm. In each step, we extend each text in the batch by one character. In the following, $v$ and $w$ are the current and the next character in the current text $T_j$ and $i$ is a record offset. If $v$ is the last character of the text (the endmarker is at $T_j[0]$), we set $w = \$$. In each step, we:

1. **Rebuild records:** The texts are sorted by positions $(v, i)$ such that the endmarker of that text should be at $\mathsf{BWT}_v[i]$. (We do not write the temporary endmarkers to the records.) We process all texts at the same node $v$ to rebuild the record.

   (a) If the record does not contain the edge $(v, w)$, we add $(w, 0)$ to the header.

   (b) We add BWT runs and text identifiers until offset $i$ to the new record. If we have inserted $k$ characters so far, we replace text identifier $(i', j')$ with $(i' + k, j')$.

   (c) If $w = \$$ or the text position is divisible by $d$, we insert text identifier $(i, j)$.

   (d) We insert $w$ to the BWT and set $i \leftarrow \mathsf{BWT}_v.\mathsf{rank}(i, w)$.

   (e) If $w \neq \$$, we increment the number of paths from $v$ to $w$ in the incoming edges of $w$.

2. **Sort:** We sort the texts by $(w, v, i)$, which is the order we need in the next step. If $w = \$$, the text is now fully inserted, and we remove it from further processing.

3. **Rebuild offsets:** For each distinct node $w$, we rebuild the $\mathsf{BWT.rank}(v', w)$ fields in the outgoing edges of predecessor nodes $v'$ using the path counts in the incoming edges of $w$. Then we set $i \leftarrow i + \mathsf{BWT.rank}(v, w)$ to have the correct offset in the next step.

#### 2.2 GBWT merging

The GBWT construction algorithm is sequential. Parallelizing it is difficult, because the algorithm interleaves queries with index updates. For faster construction, we can build GBWT indexes for multiple batches in parallel and then merge the partial indexes.

Let $\mathsf{GBWT}_1$ and $\mathsf{GBWT}_2$ be two GBWT indexes we want to merge. The basic idea [6] is the following. We extract texts from $\mathsf{GBWT}_2$ and search for them in $\mathsf{GBWT}_1$ as in direct construction. Instead of updating $\mathsf{GBWT}_1$, we only store the positions $(v, i)$ where we would have updated $\mathsf{GBWT}_1$. The sorted positions form the *rank array* of $\mathsf{GBWT}_2$ relative to $\mathsf{GBWT}_1$. The rank array tells us how we can build the merged index by interleaving $\mathsf{GBWT}_1$ and $\mathsf{GBWT}_2$.

Our GBWT merging algorithm is based on BWT-merge [7]. In the *search phase*, we extract texts from the dynamic $\mathsf{GBWT}_2$ and search for them in dynamic $\mathsf{GBWT}_1$ using multiple search threads. Because we expect that the texts are long and that most of their suffixes are unique, each thread extracts one text at a time. In contrast, the original BWT-merge was intended for merging the BWTs of short read collections. It traversed the trie of reverse texts and reported each distinct suffix only once.

Each search thread has a *position buffer* and a *thread buffer*. The reported positions are stored in the position buffer. When the buffer gets full, we sort the positions, encode them differentially, and merge the compressed buffer with the thread buffer. (The original BWT-merge had run-length encoded buffers, because the search threads reported multiple occurrences of the same suffix.) If the thread buffer also gets full, we try to insert it into the global *merge buffers*, starting from buffer 0. Each merge buffer $i$ is either empty or contains $2^i$ merged thread buffers. If the merge buffer $i$ we are currently trying is empty, we insert the thread buffer into it and return to searching. Otherwise we take the existing merge buffer, merge it with the thread buffer, and continue to buffer $i + 1$. If there are no more merge buffers remaining, we write the thread buffer to a new file.

In the *merge phase*, we use the rank array for interleaving the GBWTs. There is a separate *reader thread* for reading and decompressing each file. The *merge thread* takes the streams generated by the reader threads and merges them into a single stream using a tournament tree. (There was no separate merge thread in the original BWT-merge.) The *main thread* takes that stream, interleaves the BWTs, and updates the stored text identifiers.

Because searching in a dynamic GBWT is fast, the multithreaded search phase can be much faster than the effectively sequential merge phase. We use multiple *merge jobs* to avoid this sequential bottleneck, which did not exist in the original BWT-merge. We partition the alphabet between the merge jobs, so that each job gets a range $[a, b] \subseteq \Sigma$. When a search thread writes its thread buffer to disk, it creates a separate file for each partition. In the merge phase, each merge job has a separate rank array, which it uses for interleaving the range of records $[a, b]$.

When the node identifiers in $\mathsf{GBWT}_1$ and $\mathsf{GBWT}_2$ do not overlap (e.g. we have indexes for different chromosomes), merging is much faster. We can also merge more than two indexes in this case. The records for all nodes $v \in V$ can be reused in the merged index. In the endmarker record \$, we merge the local alphabets and concatenate the record bodies. We also have to update the stored text identifiers in all records, assigning new identifiers according to the order we used in the endmarker.

Texts can be removed from the GBWT by reversing the merging algorithm. We extract the texts we want to remove and store the positions we encounter. Once we have the rank array, we

| Dataset | Index | find($X$), $\lvert X\rvert = 2$ Unidir | find($X$), $\lvert X\rvert = 2$ Bidir | find($X$), $\lvert X\rvert = 50$ Unidir | find($X$), $\lvert X\rvert = 50$ Bidir | locate() Queries | locate() Length | locate() Direct | locate() Fast | extract() Length | extract() Time |
|---|---|---|---|---|---|---|---|---|---|---|---|
| 1000GP-all-S | Compressed | 460 ns | 530 ns | 220 ns | 260 ns | 20,000 | 57.1 M | 96 µs | 11 µs | 433 M | 160 ns |
| 1000GP-all-S | Dynamic | 150 ns | 260 ns | 74 ns | 95 ns | 20,000 | 57.1 M | 19 µs | 9.4 µs | 433 M | 790 ns |
| 1000GP-all-L | Compressed | 470 ns | 540 ns | 220 ns | 260 ns | 20,000 | 57.1 M | 110 µs | 11 µs | 91.8 G | 170 ns |
| 1000GP-all-L | Dynamic | 150 ns | 260 ns | 77 ns | 94 ns | 20,000 | 57.1 M | 19 µs | 9.6 µs | 91.8 G | 89 ns |
| TOPMed-17-L | Compressed | 400 ns | 500 ns | 260 ns | 330 ns | 200 | 13.0 M | 1600 µs | 220 µs | 216 G | 200 ns |
| TOPMed-17-L | Dynamic | 140 ns | 240 ns | 82 ns | 110 ns | 200 | 13.0 M | 360 µs | 210 µs | 216 G | 100 ns |

Table 1: Query benchmarks. find(): We give the average query time in nanoseconds/character for each dataset and index type for pattern lengths $|X| = 2$ and $|X| = 50$ in both unidirectional and bidirectional search. locate(): We give the number of query ranges, total length of the query ranges, and average time in microseconds/position with direct and fast algorithms. extract(): We give the total length of the extracted paths and the average time in nanoseconds/character. M and G suffixes denote millions and billions, respectively.

remove the marked positions from the BWT. We also have to update the text identifiers to remove the gaps we may have created. Because we usually want to remove only a small number of texts (e.g. those corresponding to a particular sample), we can store the rank array as an uncompressed array in memory.

##### 2.3 Merging the TOPMed superbatches

We merged the superbatch indexes using BWT-merge with 32 search threads, 64 MiB position buffers, 256 MiB thread buffers, 6 merge buffers, and 8 merge jobs. The rank arrays were written to disk in 1.5 GiB files, and the peak disk usage was 575 GiB. The time for merging a new superbatch into the index varied between 2.0 hours and 2.4 hours, except for the last superbatch which took 1.4 hours. Roughly 75% of the time was spent in the search phase and 25% in the merge phase. During the merge phase, merge jobs read the rank arrays at an average rate of 250 MiB/s.

#### 3 GBWT benchmarks

We benchmarked the basic queries on both whole-genome 1000GP indexes and on the chromosome 17 TOPMed index. For find(), we selected several pattern lengths $|X|$ from 2 to 50 and extracted $100,000$ patterns of each length, starting from random positions in the index. We then measured the average time per character in unidirectional and bidirectional search. This can be understood as measuring $\mathsf{O}(t_r)$ in the expected case, where $t_r$ is the time required to compute rank on the BWT. See Table 1 for the results.

As expected, the average time per character was lower with long patterns due to memory locality. Bidirectional search was slightly slower than unidirectional search. The overall smaller TOPMed chromosome 17 index was faster with short patterns than the 1000GP indexes, because random access times are lower in smaller structures. The situation was reversed with long patterns, as we benefit less from memory locality when the individual records are larger. For a comparison, the typical find() speed in uncompressed FM-indexes for DNA sequences is 200 to 400 ns/character [8], which is comparable to the compressed GBWT. The dynamic GBWT is several times faster.

The locate() performance suffers from the long distance between stored identifiers: $d = 1,024$ in the 1000GP indexes and $d = 16,384$ in the TOPMed index. We extracted a number of patterns of length 20 from the index and used the ranges returned by $\mathsf{find}()$ queries for $\mathsf{locate}()$ benchmarks. We measured the average time per occurrence. With the direct algorithm, this corresponds to measuring $\mathsf{O}(d \cdot t_r)$. See Table 1 for the results.

With the direct $\mathsf{locate}()$ algorithm, the dynamic index was several times faster than the compressed index. The fast algorithm improved the performance of the compressed index by an order of magnitude, making the difference between the index types minimal. Queries were 16 to 22 times slower in the TOPMed index than in the 1000GP indexes due to the longer distance between stored identifiers. For a comparison, FM-indexes for non-repetitive text typically use $d = 16$ or $d = 32$ and take a few microseconds to locate each position [8].

We also extracted 10,000 paths from each index and measured the average time per character. This corresponds to measuring $\mathsf{O}(t_r)$. See Table 1 for the results. The $\mathsf{extract}()$ times are comparable to long $\mathsf{find}()$ queries, except for the dynamic index for short 1000GP paths. While the compressed index stores the decompressed endmarker record and uses it for $\mathsf{extract}()$ queries, we do not do this in the dynamic index, as we expect the index to change frequently. When the total number of paths is large and the extracted paths are short, the majority of the time is spent decompressing the endmarker. (When we extract paths from the dynamic index in the BWT-merge algorithm, we decompress the endmarker only once.)

### 4 Input and output sizes

| | Input | | | Output | | |
|---|---|---|---|---|---|---|
| **Dataset** | **Reference** | **VCF** | **Total** | **Graph** | **GBWT** | **Total** |
| 1000GP-all-S | 838 MiB | 16.2 GiB | 17.0 GiB | 4.14 GiB | 18.6 GiB | 22.8 GiB |
| 1000GP-all-L | 838 MiB | 16.2 GiB | 17.0 GiB | 4.14 GiB | 16.6 GiB | 20.8 GiB |
| 1000GP-17-S | 22 MiB | 414 MiB | 437 MiB | 115 MiB | 500 MiB | 615 MiB |
| 1000GP-17-L | 22 MiB | 414 MiB | 437 MiB | 115 MiB | 444 MiB | 559 MiB |
| TOPMed-17-L | 25 MiB | 9.16 GiB | 9.18 GiB | 329 MiB | 2.16 GiB | 2.48 GiB |

Table 2: Input and output sizes for the datasets. The input consists of gzip-compressed reference and VCF files, while the output consists of VG graph and GBWT index files.
